# Phylogenetic clustering of the Indian SARS-CoV-2 genomes reveals the presence of distinct clades of viral haplotypes among states

**DOI:** 10.1101/2020.05.28.122143

**Authors:** Bornali Bhattacharjee, Bhaswati Pandit

**Author notes:** **Correspondance: Bornali Bhattacharjee, Bhaswati Pandit**, National Institute of Biomedical Genomics, Kalyani, West Bengal, India.

## Abstract

The first Indian cases of COVID-19 caused by SARS-Cov-2 were reported in February 29, 2020 with a history of travel from Wuhan, China and so far above 4500 deaths have been attributed to this pandemic. The objectives of this study were to characterize Indian SARS-CoV-2 genome-wide nucleotide variations, trace ancestries using phylogenetic networks and correlate state-wise distribution of viral haplotypes with differences in mortality rates. A total of 305 whole genome sequences from 19 Indian states were downloaded from GISAID. Sequences were aligned using the ancestral Wuhan-Hu genome sequence (NC_045512.2). A total of 633 variants resulting in 388 amino acid substitutions were identified. Allele frequency spectrum, and nucleotide diversity (π) values revealed the presence of higher proportions of low frequency variants and negative Tajima’s D values across ORFs indicated the presence of population expansion. Network analysis highlighted the presence of two major clusters of viral haplotypes, namely, clade G with the S:D614G, RdRp: P323L variants and a variant of clade L [L_v_] having the RdRp:A97V variant. Clade G genomes were found to be evolving more rapidly into multiple sub-clusters including clade GH and GR and were also found in higher proportions in three states with highest mortality rates namely, Gujarat, Madhya Pradesh and West Bengal.

## 1. Introduction

Coronavirus disease 2019 (COVID-19) caused by the Severe Acute Respiratory Syndrome Coronavirus 2 (SARS-CoV-2) was first reported in December, 2019 from Wuhan, China and since then has spread across the globe with 349,190 deaths as reported to WHO[1, 2]. The SARS-CoV-2 has a 29.9 kilobase long RNA genome coding for 22 proteins. The first whole genome (RNA) sequence of SARS-CoV-2 was published on the 5^th^ of January, 2020 [3] and currently more than 30,000 SARS-CoV-2 sequences have been submitted from across the world to Global Initiative on Sharing All Influenza Data (GISAID)[4]. It has also been identified on the basis of nucleotide variants that 8 major clades of viral haplotypes have spread across the globe causing the pandemic [4]. However, the implications of the evolutionary genome-wide changes still remain elusive.

Sequencing of SARS-CoV-2 is imperative to understand the transmission routes, possible sources and cross species evolution and transmission to human hosts. On the basis of such sequence identity it has been speculated that the bats form reservoir of such viruses (bat CoV genome, RaTG13) and are a probable species of origin [5]. Further, reports have also shown strong homology among viruses in metavirome data sets of SARS-CoV, which were generated from the lungs of deceased pangolins [6].

In India, the first three cases of COVID-19 with travel history from Wuhan, China were reported from the state of Kerala in February 2020. Since then the virus and the disease has spread to all 37 states and union territories with 86110 active cases and 4531 deaths till date and the percentage of death rates seem to differ among states so far [7]. Attempts have been made to sequence the genomes of Indian clinical isolates to understand genome-wide variability and viral evolution and over 300 sequences have been deposited to GISAID so far from many Indian states [8, 9]. However, there has been no study to delineate ancestries or to characterize the distribution patterns of viral haplotypes across states. Hence, in this study a total of 305 Indian SARS-CoV genome sequences were used in an effort to understand the evolution of these viruses, trace the routes of infection and gauge the clustering patterns across states.

## 2. Material and Methods

### 2.1 Nucleotide alignment, and variant calling

All SARS-CoV-2 genome sequences (n=305) that had been isolated from Indian nationals and submitted from India were downloaded on the 18^th^ of May 2020 from the GISAID database. Out of a total of 305 sequences, 26 were found without state information and 7 had been grown in Vero cells. The rest of the isolates were collected from 20 different states which included Andhra Pradesh (n=2), Assam (n=2), Bihar (n=6), Delhi (n=39), Gujarat (n=103), Haryana (n=1), Jammu (n=1), Kashmir (n=1), Karnataka (n=17), Kerela (n=2), Ladakh (n=6), Madhya Pradesh (n=10), Maharastra (n=7), Odisha (n=1), Punjab (n=1), Rajasthan (n=1), Tamil Nadu (n=19), Telangana (n=35), Uttar Pradesh (n=5) and West Bengal (n=13) (Supplementary table S1). Given the initial emergence of the SARS-CoV-2 virus from Wuhan, China, alignment was carried out using the genome sequence submitted by Wu *et. al.* (NC_045512.2) in January 2020[3]. Additionally, the bat and pangolin SARS-related coronaviruses [RaTG13 (MN996532), MN789 (MT121216)] that have been reported to be the closest to the SARS-CoV-2 virus [10] were aligned to identify conserved amino acids across ORFs. Multiple sequence alignment was executed using MUSCLE [11] with three iterations for both. Since different sequencing platforms had been used to generate SARS-CoV-2 sequence data with different error rates and filtering cut-offs hence, the variant calling was stringently carried out with a minimum coverage of 150 genomes. Ambiguous bases found in genome sequences were considered to be unresolved for the purpose of analyses. The SIFT database was used to identify amino acid changes that could protein function (http://blocks.fhcrc.org/sift/SIFT_seq_submit2.html) [12].

### 2.2 Measurements of diversity and deviation from neutrality

Watterson’s estimator (θ_w_), nucleotide diversity (π) and Tajima’s D [13] for each open reading frame (ORF) was calculated using MEGA X [14].

### 2.3 Phylogeny construction

Phylogenetic analysis was carried out following the median-joining approach using Network 10.1.0.0 software [15]. For phylogenetic construction a variant frequency cutoff of ≥ 0.01 was used and a 97% cutoff for the number of sequences with resolved bases for each position to avoid spurious clustering. One genome sequence (EPI_ISL_414515) had to be removed from analyses for the absence of complete sequence information.

## 3. Results

### 3.1 Description of the variants found within the Indian SARS-CoV-2 genomes

Multiple sequence alignment with 305 Indian viral sequences and ancestral Wuhan-Hu-1 isolate sequence (NC_045512.2) revealed the presence of 572 single nucleotide variants along with 61 di-nucleotide, tri-nucleotide substitutions, insertions and deletions out of which 388 were non-synonymous changes resulting in amino acid substitution. The allele frequencies of all the 633 variants varied from 0.3-53.4% and greater than 90% of these were low frequency variants ranging from 0.3-1% (Figure 1A, supplementary table S2). Among the single nucleotide variants C-T transition per cytosine residue in the genome was highest at 3.04%, followed by G-T transversion (1.74%) (Figure 1B, supplementary table 2). The NSP3 and S ORFs had the highest number of non-synonymous changes each (n=75, 19.33%) (Figure 1C).

**Figure 1:**
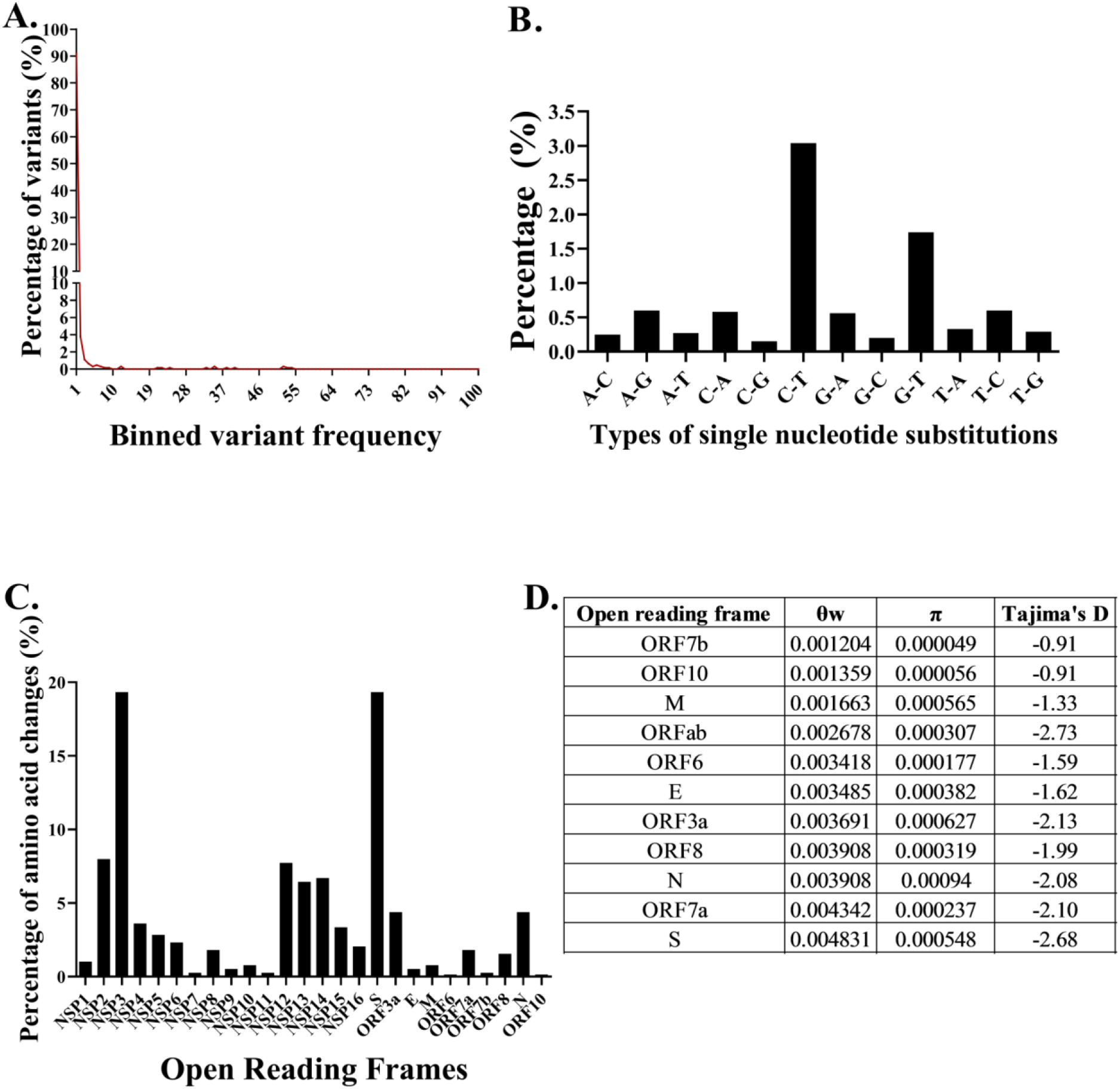
Description of SARS-CoV2 variants, the gene diversities and the results of the test for neutrality. A. The frequencies of variants binned on the basis of frequency. B. The types of single nucleotide changes incurred by the viral genomes C. The percentage of amino acid changes per open reading frame out of a total of 388 non-synonymous changes that were observed. D. The diversity and Tajima’s D values across open reading frames.

### 3.2 Deviation from neutral viral evolution

Given the number of variants identified, the diversities across all the ORFs were calculated. ORF7a was found to have the lowest nucleotide diversity while the S ORF had the highest. Overall, the nucleotide diversity (π) values were low across ORFs in comparison to the θ (Watterson’s estimater) values (Figure 1D) which was indicative of the presence of higher proportion of low frequency variants as has been described in Figure 1A. The next objective was to determine if the patterns of diversity could be attributed to genetic drift or neutrality. Tajima’s test for neutrality was applied and all the ORFs were found to have negative Tajima’s D values (Figure 1D) indicative of non-neutral evolution.

### 3.4 Presence of two major clades of Indian SARS-CoV-2 haplotypes with emergence of multiple subclusters

To trace the ancestries of the Indian viral isolates, network analysis was carried out using the Hamming distances of variants present at ≥1% frequency among the genomes (Supplementary table S3). The haplotypes were generated stringently using a total of 53 single nucleotide variants, a di-nucleotide variant AA-CU at positions 10,478-10,479, a tri-nucleotide substitution at positions 28,881-28,883GGG-AAC together resulting in 38 amino acid changes and 304 Indian SARS-CoV-2 genome sequences (Figure 2, Supplementary table S4). A total of 54 nodes with 54 distinct haplotypes were discovered using 304 genome sequences. The network appearance was as expected from an ongoing pandemic with the presence of ancestral viral haplotypes existing along with newly mutated genomes. There were two major clusters of haplotypes that were found to have emerged from the ancestral Wuhan-Hu-1 virus (clade L) identified to be belonging to clade G and a variant of the clade L which has been annotated here as L_v_ (Figure 3). Seventy-nine viral isolates were found to be belonging to clade L_v_ and there were 36 clade G viruses. The clade L_v_ genomes differed from the ancestral Wuhan-Hu-1 sequence at one locus resulting in a change in the RNA dependent RNA polymerase (RdRp) protein [C13,730U,(A97V)] while the G haplotype viruses differed at three loci resulting in one amino acid change each in the RdRp, S protein [C3037U, C14,408U (RdRp: P323L), A23,403G (S:D614G)]. Two sub-clusters were observed evolving from clade G; GH variant mentioned here as GH_v_ with the variants C18877U, G25563U (ORF3a:Q57H), C26735U having multiple evolving branches and GR with the tri-nucleotide GGG-AAC substitution at positions 28,881-28,883 resulting in two amino acid changes R203K, G204R in the N protein (Figure 3). Reticulations were also observed which could happen because of parallel mutations or homoplasy.

**Figure 1:**
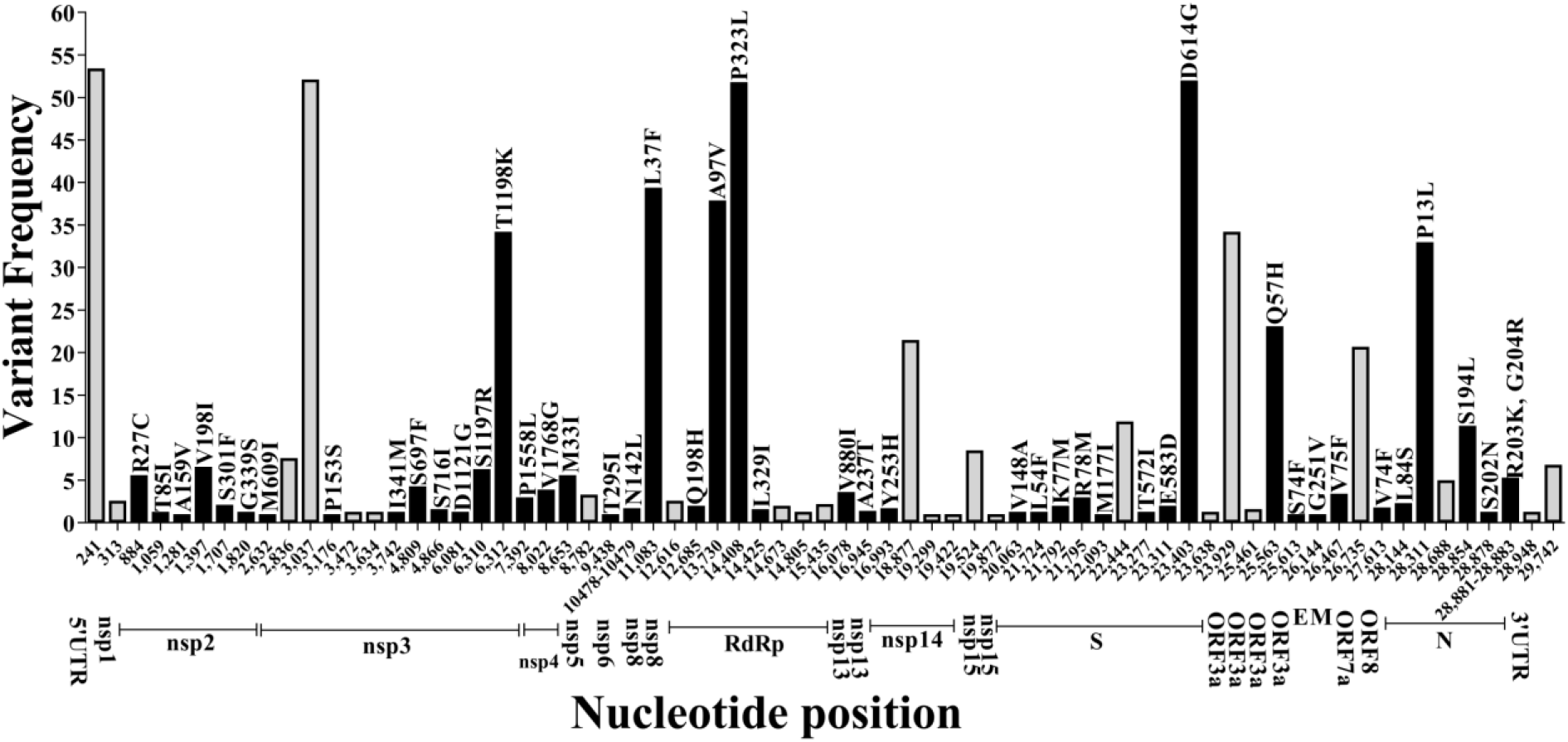
Distribution of nucleotide variants on the basis of which ancestries were derieved with variant frequencies and associated amino acid changes. The black bars represent the non-synonymous variants with the amino acid changes mentioned above and the coding regions are indicated below.

**Figure 3:**
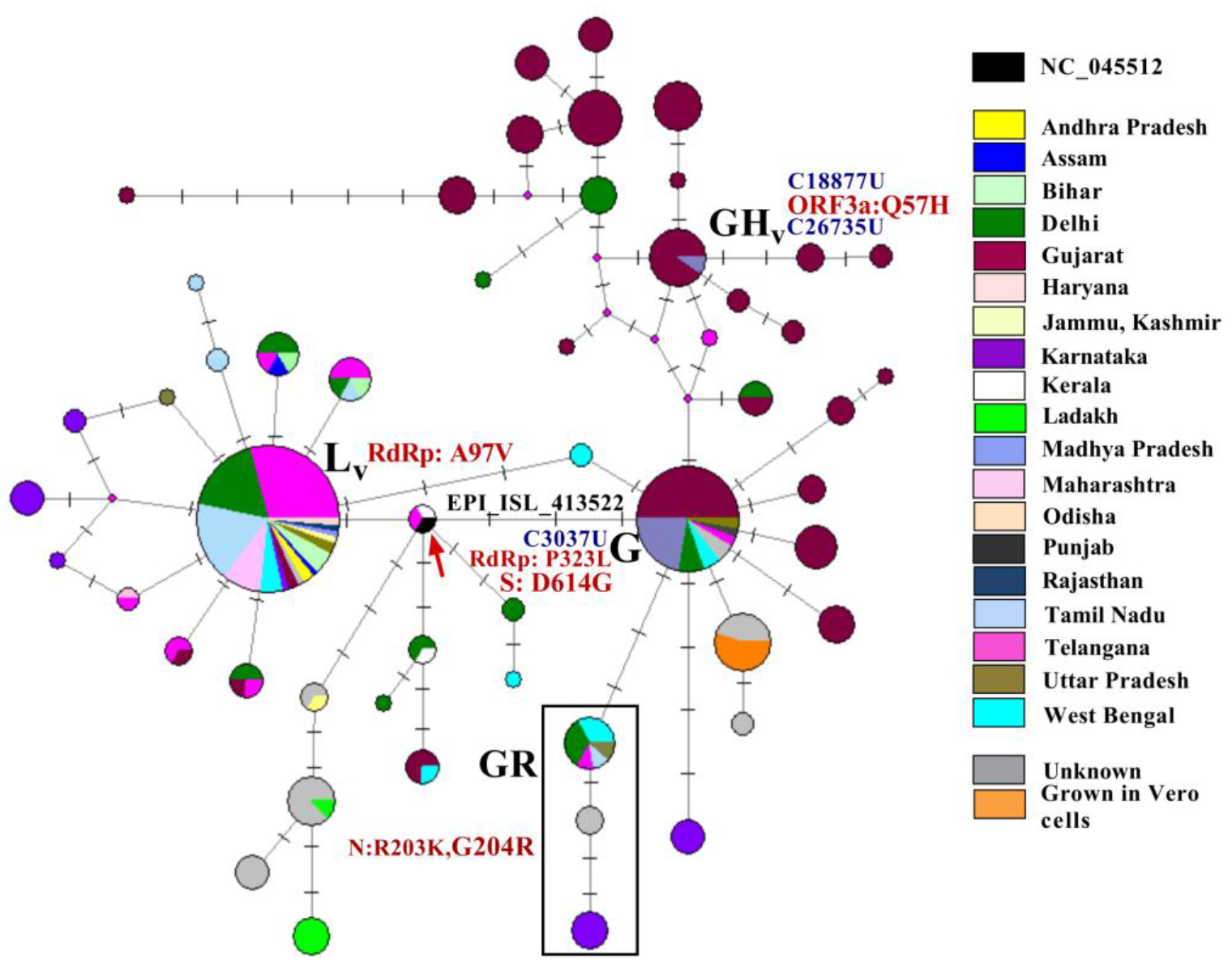
Phylogenetic network of 304 Indian SARS-CoV-2 viral genomes. Each circle represents a haplotype and the diameter is proportional to the number of genomes belonging to each haplotype. Each notch on a horizontal line represent a differentiating variant and the length of the connecting lines are proportional to the number of variants. The colors indicate the different states from which specimens were collected. The arrow indicates the node containing the ancestral NC_0145512 viral haplotype and the sequence ID indicate the first sequenced viral genome from a symptomatic individual with travel history to Wuhan, China. A total of 55 variants were used to construct the haplotypes and the nucleotide and amino acid differences in comparison with NC_045515 among the major clades L_v_(RdRp:A97V), G, GH_v_[C18887U, C26735U] and GR with the 48881-48883(GGG-AAC, N:R203K, G204R) variants are mentioned.

There were 14 variants [C241U, C1707U, G2632U, C6310A, C6312A, U8022G, G11083U, A15435G, U19299G, A19422G, C19524U, C23929U, G27613U, C28311U] that could not be included in the network analysis because of the presence of unresolved bases at multiple viral genome sequences. These variants were subsequently evaluated in the subset of sequences that had information among the major haplotypes belonging to clade L_v_ (n=28) and G (n=20). The variants G11083U (NSP6:L37F), C23929U, C28311U (N:P13L) were found to cluster in high proportion of clade L_v_ viruses (71.5%) and as expected the 5’ UTR variant C241T was found to cluster among the clade G viral isolates and its evolving sub-clusters.

All the amino acid changes that were present at ≥1% frequency were also evaluated on the basis of conservation (Table 1). NSP3 amino acid changes were found to be at the most non-conserved sites; however, the changes were predicted to be affecting protein function. Further, there were three loci where the variants resulted in amino acid changes that were fixed in either the RaTG13 or the MN789 genome (NSP2: V198I, NSP3: D1121G, T1198K). Most of the variants across clades L_v_, G, GH_v_ and GR were found to occur in conserved codons across species except for the NSP6: L37F variant where the leucine codon had been replaced by a valine in the MN789 genome and the S:D614G variant where the pangolin coronavirus isolate MN789 S-ORF had a threonine codon.

**Table 1:**
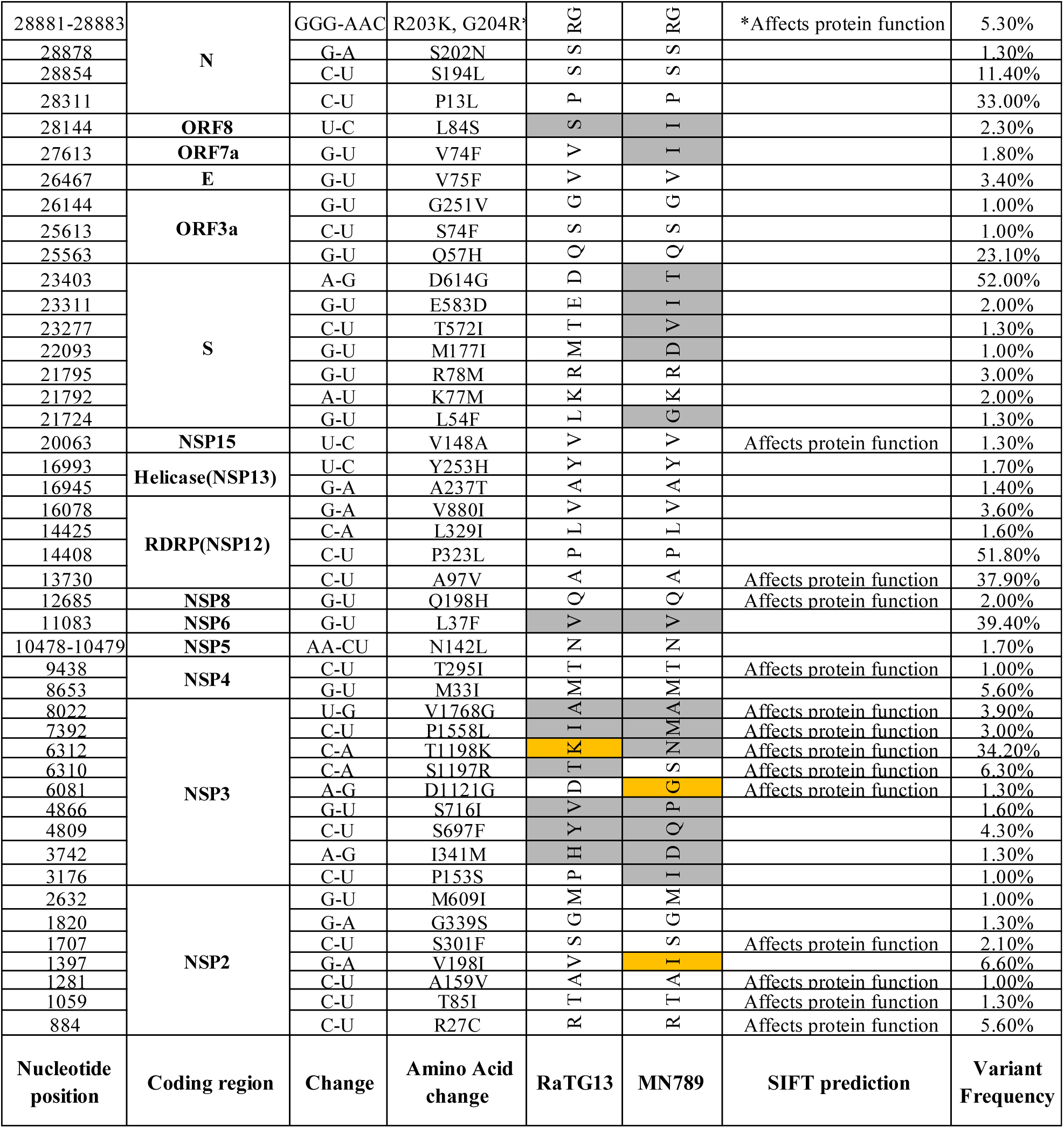
The amino acid changes present at ≥1% frequency in each ORF across conserved and non-conserved positions in different ORFs. The protein function affecting changes are marked according to SIFT scores. The grey cells indicate non-conserved positions and the yellow cells indicate variants that code for the same amino acids as fixed either in bat coronavirus RaTG13 or pangolin coronavirus MN789 genomes.

### 3.4 Statewise distribution of viral haplotypes

There were two individuals with whole genome sequence data (sequence IDs EPI_ISL_413522, EPI_ISL_413523) from Kerala who were the first to be identified with COVID-19 symptoms and specimens were collected from India on 27^th^ January and 31^st^ January 2020 respectively [9]. It was found that the EPI_ISL_413522 haplotype clustered with the Wuhan-Hu-1 haplotype and the second isolate had two nucleotide changes resulting in an amino acid change in the ORF8 protein [C8782U, U28,144C (ORF8: L84S)] (Figure 3). Additionally, a viral isolate collected on 1^st^ March 2020 from Telangana was also found to cluster with the ancestral isolate.

So far the highest number of deaths per number of confirmed cases has been recorded in the states of Gujarat (6.2%), Madhya Pradesh (4.3) and West Bengal (6.9%) (Supplementary table S5). Out of these three states the maximum numbers of isolates have been sequenced from Gujarat (n=103). Among the total of 103 of these viral isolates, 18 (17.5%) were found to belong to clade G, while the rest with the exception of 7 L_v_ clade isolates (6.8%) were distributed among sub-clusters arising from clade G (n=77; 74.8%) with additional number of variants. Among the 7 clade L_v_ viral isolates, 6 were observed to have been isolated from individuals of the same city, Surat. The isolates from Madhya Pradesh (n=10) were also distributed between clade G and its sub-cluster, clade GH_v_ while >50% (7 out of 13) the West Bengal isolates belonged to clade G, GR or another sub-cluster of clade G with the S:D614G variant. The viral isolates(n=6) from Bihar which has much lower mortality rates (0.5%) were found to be distributed among the clade L_v_ and its branches and the isolates from Ladakh (n=6) which has not had any COVID-19 deaths were found to differ from the ancestral isolate with three non-synonymous changes [C884U (nsp2: R27C), G1397A (NSP2: V198I), G8653U (NSP4: M33I)], a synonymous change (U28688C), a change in the 3’UTR region [G29742U] and one non-synonymous and another synonymous change in 5 of the 6 isolates [U16993C (nsp13: Y253H), U25461C]. The isolate sequences from Delhi which has 2% mortality were found to be distributed across the both the major clusters and sub-clusters.

## 4. Discussion

A number of studies from India have described the presence of various mutations across the SAR-CoV-2 genome with hints of selection and *in silico* predictions of alterations in protein functions [16–18] so far. In this study an attempt has been made to understand the pattern of evolution within human hosts, to infer the ancestries of the viral isolates using the phylogenetic network approach which had been used earlier to build SARS-CoV-2 infection paths [19] and make an effort at correlating it with morbidity.

While comparing the 305 Indian SARS-CoV-2 genomes a number of nucleotide variants or segregating sites were identified, however, the nucleotide diversity values (π) were indicative of an excess of low frequency variants. This could be because of recent population expansions as has been observed in H1N1 populations involved in outbreaks and epidemics [20] and the uniform negative Tajima’s D values across all the SARS-CoV-2 ORFs could also be attributed to it.

The clade G (S:D614G) viruses with the C14,408U (RdRp: P323L) variant were found to incur more number of mutations leading to the emergence of a number of sub-clusters of viruses with increased branch lengths in comparison to the clade L_v_ viruses including clade GH and GR. These clade G viruses were also found in more numbers in states where higher mortality rates were recorded. Occurrence of higher numbers of mutations might be attributed to altered secondary structure and impaired RdRp proofreading due to the C14,408U (RdRp: P323L) variant as has been speculated in earlier reports [21, 22]. Additionally, there has been a report on clinical outcome from Sheffield, England where the G614 mutation was associated with higher viral loads [23] which might be contributing to the higher mortality rates. However, these implications will have to be tested further with direct correlations using comprehensive clinical data and genomic data from all the states.

## Supplementary materials

S1 table: Details of all the sequences included in the study.

S2 table: Description all the variants identified in this study.

S3 table: Features of the variants included in the construction of viral haplotypes.

S4 table: Haplotype information.

S5 table: Official state-wise numbers of confirmed COVID-19 cases and deaths from 19 Indian states as updated on 28^th^ May 2020.

## Acknowledgments

The authors acknowledge the submitters of coronavirus sequence data to the GISAID database, the database managers, developers and scientists associated with GISAID and Prof. Saumitra Das for encouragement.

## Author Contributions

Conceptualization, B.B.; data curation, B.B.; data analysis, writing-original draft, B.B. & B.P.; writing-review and editing, B.B. & B.P. Both authors approved the manuscript.

## Competing interests

None declared.

## Funding

B.B was supported by the Ramanujan fellowship funded by the Department of Science and Technology, Government of India.

## Notes

### Competing Interest Statement

The authors have declared no competing interest.

## References

1. Lu, R.; Zhao, X.; Li, J.; Niu, P.; Yang, B.; Wu, H.; Wang, W.; Song, H.; Huang, B.; Zhu, N.; Bi, Y.; Ma, X.; Zhan, F.; Wang, L.; Hu, T.; Zhou, H.; Hu, Z.; Zhou, W.; Zhao, L.; Chen, J.; Meng, Y.; Wang, J.; Lin, Y.; Yuan, J.; Xie, Z.; Ma, J.; Liu, W. J.; Wang, D.; Xu, W.; Holmes, E. C.; Gao, G. F.; Wu, G.; Chen, W.; Shi, W.; Tan, W., Genomic characterisation and epidemiology of 2019 novel coronavirus: implications for virus origins and receptor binding. Lancet 2020, 395, (10224), 565–574.

2. (https://covid19.who.int/). WHO 2020.

3. Wu, F., Zhao, S., Yu, B. et al., A new coronavirus associated with human respiratory disease in China. Nature 2020, 579, 265–269

4. https://www.gisaid.org/. 2020.

5. Zhou, H.; Chen, X.; Hu, T.; Li, J.; Song, H.; Liu, Y.; Wang, P.; Liu, D.; Yang, J.; Holmes, E. C.; Hughes, A. C.; Bi, Y.; Shi, W., A Novel Bat Coronavirus Closely Related to SARS-CoV-2 Contains Natural Insertions at the S1/S2 Cleavage Site of the Spike Protein. Current biology: 2020.

6. Liu, P.; Chen, W.; Chen,P.; Viral Metagenomics Revealed Sendai Virus and Coronavirus Infection of Malayan Pangolins (Manis javanica). Viruses 2019, 11, (11), 979.

7. https://www.mygov.in/corona-data/covid19-statewise-status/. 2020.

8. Maitra, A.; Chawla Sarkar., M.; Rajeja, H.; Biswas N.K.; Chakraborti, S.; Singh,A.K.; Ghosh, S.; Sarkar, S.; Patra, S.; Mandal, R.K.; Ghosh,T. et.al., Mutations in SARS Cov2 viral RNA identified in Eastern India: Possible implication for the ongoing outbreak in India and impact on viral structure and host susceptibility. J Biosciences 2020, 45.

9. Yadav, P. D.; Potdar, V. A.; Choudhary, M. L.; Nyayanit, D. A.; Agrawal, M.; Jadhav, S. M.; Majumdar, T. D.; Shete-Aich, A.; Basu, A.; Abraham, P.; Cherian, S. S., Full-genome sequences of the first two SARS-CoV-2 viruses from India. The Indian journal of medical research 2020, 151, (2 & 3), 200–209.

10. Tang, X. Wu., C.; Li, X.;Song, Y.; Yao,X.; Wu, X.; Duan, Y.; Zhang, H.; Wang, Y.; Qian,Z, On the origin and continuing evolution of SARS-CoV-2 National Science Review 2020.

11. Edger, R. C., MUSCLE: multiple sequence alignment with high accuracy and highthroughput Nucleic Acids Research 2004, 32, (5), 1792–1797.

12. Ng, P. C.; Henikoff., S, Predicting Deleterious Amino Acid Substitutions. Genome Research 2001, 11(5):863–74.

13. Tajima, F., Statistical methods to test for nucleotide mutation hypothesis by DNA polymorphism. Genetics 1989, 123, 585–595.

14. Kumar, S.; Stecher G.; Li M.; Knyaz C.; Tamura, K. MEGA X: Molecular Evolutionary Genetics Analysis across computing platforms. Molecular biology and evolution 2018, 35, 1547–1549.

15. Bandelt H-J, Forster, P., Röhl A. Median-joining networks for inferring intraspecific phylogenies. Mol. Biol. Evol. 1999, 16, 37–48.

16. Banerjee, A.; Sarkar, R.; Mitra,S.; Mahadeb Lo, M.; Dutta, S.; Chawla-Sarkar,M., The novel Coronavirus enigma: Phylogeny and mutation analyses of SARS-CoV-2 viruses circulating in India during early 2020. bioRxiv 2020, 2020.05.25 114199.

17. Begum, F.; Mukherjee, D.; Thagriki, D.; Das, S.; Tripathi,P.P.; Banerjee,A.K.; Ray, U., Analyses of spike protein from first deposited sequences of SARS-CoV2 from West Bengal, India. bioRxiv 2020, 2020.04.28.066985.

18. Bhowmik, D.; Pal, S.; Lahiri, A.; Talukdar, A.; Paul, S., Emergence of multiple variants of SARS-CoV-2 with signature structural changes. bioRxiv 2020, 2020.04.26.062471.

19. Forster, P.; Forster, L.; Renfrew,C.; Forster,M. Phylogenetic network analysis of SARS-CoV-2 genomes. PNAS 2020, 117, (17), 9241–9243.

20. Martinez-Hernandez, F.; Jimenez-Gonzalez, D. E.; Martinez-Flores, A.;Villalobos-Castillejos, G.;Vaughan, G.;Kawa-Karasik, S.;Flisser, A.;Maravilla, P.;Romero-Valdovinos, M., What happened after the initial global spread of pandemic human influenza virus A (H1N1)? A population genetics approach. Virology journal 2010, 7, 196.

21. Pachetti, M.; Marini, B.; Benedetti, F.; Giudici, F.; Mauro, E.; Storici, P.; Masciovecchio, C.; Angeletti, S.; Ciccozzi, M.; Gallo, R. C.; Zella, D.; Ippodrino, R., Emerging SARS-CoV-2 mutation hot spots include a novel RNA-dependent-RNA polymerase variant. Journal of translational medicine 2020, 18, (1), 179.

22. Chand, G. B.; Banerjee, A.; Azad G.K. Identification of novel mutations in RNA-dependent RNA polymerases of SARS-CoV-2 and their implications on its protein structure. bioRxiv 2020, 2020.05.05.079939.

23. Korber, B.; Fischer. W., Gnanakaran S, Yoon H, Theiler J, Abfalterer W, Foley B, Giorgi EE, Bhattacharya T, Parker MD, Partridge DG, Evans CM, de Silva T, on behalf of the Sheffield COVID-19 Genomics Group, LaBranche CC,; Dc, M., Spike mutation pipeline reveals the emergence of a more transmissible form of SARS-CoV-2. bioRxiv 2020, 2020.04.29.069054.

